# Cerebellar systems consolidation driven by the temporal dynamics of Purkinje cell excitability

**DOI:** 10.1101/2024.07.24.604888

**Authors:** Jewoo Seo, Yong Gyu Kim, Yong-Seok Lee, Sang Jeong Kim

## Abstract

Systems consolidation, essential for long-term memory formation, orchestrates the reorganization of newly encoded memories from the cortex into subsequent neural circuitry. While the role of synaptic mechanism in consolidation is well understood, the contribution of neuronal intrinsic excitability (IE) remains relatively unexplored. Herein, we adopted the optokinetic reflex, a cerebellum-dependent learning model, and manipulated IE of the sole output of the cerebellar cortex, Purkinje cells (PCs), to corroborate the direct causality between neuronal IE and memory consolidation. Optogenetic modulation of PC-IE post-learning uncovered a critical temporal window for its role in systems consolidation. Specifically, increasing PC-IE within 90 minutes after learning disrupted memory consolidation, while outside this window, memory retention remained unaffected. Under physiological conditions, PC-IE naturally decreased during this time frame and returned to baseline thereafter. Moreover, abnormally heightened PC-IE eliminated intrinsic plasticity in flocculus-targeting neurons (FTNs) within the medial vestibular nucleus (MVN), a key downstream circuit involved in long-term memory storage. These findings highlight the precise temporal dynamics of IE as a pivotal mechanism in systems consolidation, emphasizing its vulnerability to disruptions.

## Introduction

Systems memory consolidation is the process by which memory becomes stable and permanent over time (Wiltgen & Tanaka, 2013). Memories initially encoded within the hippocampal entorhinal cortex are consolidated in the neocortex for long-term storage (Kitamura et al., 2017). In the cerebellum, short-term memories initially acquired through motor learning, driven by sensory inputs from the external environment, undergo maturation and consolidation (Attwell et al., 2002; Boyden et al., 2004; De Zeeuw et al., 2021). This process involves neural communication between GABAergic inhibitory Purkinje cells (PCs) (Kayakabe et al., 2014) in the flocculus, a lobe-like structure responsible for the ocular reflex (Chang et al., 2022; Inoshita & Hirano, 2018; Okamoto et al., 2011; Shutoh et al., 2006), and flocculus-targeting neurons (FTNs) in the medial vestibular nuclei (MVN) (Sekirnjak & du Lac, 2006; Sekirnjak et al., 2003; Shin et al., 2011; Yamazaki et al., 2015). Cerebellar systems consolidation accounts for the enhancement of compensatory eye movements, such as the optokinetic reflex (OKR) (Cahill & Nathans, 2008; Schweigart et al., 1997), which relies on various sensory modalities, including visual inputs, to ameliorate the stability of vision (Lisberger, 2009).

The canonical view of cerebellar learning and memory has long been centered on bidirectional modifications in synaptic transmission, particularly long-term potentiation, and long-term depression, which are widely recognized as pivotal processes underlying memory formation and retention (De Zeeuw et al., 1998; Hansel et al., 2006; Kandel et al., 2014; Medina et al., 2000). Empirical pieces of evidence suggest that the attenuation of synaptic strength between parallel fibers and PCs plays a prominent role in the consolidation of motor memory, allowing for the adaptation of cerebellum-dependent behaviors (De Zeeuw et al., 1998; Hansel et al., 2006; Hirano, 2013; Ito, 2013).

However, a growing body of evidence has demonstrated that understanding of memory consolidation extends beyond synaptic mechanisms and involves complementary non-synaptic mechanisms, such as intrinsic excitability (IE), to achieve intact learning capabilities (Chen et al., 2020; Kim & Linden, 2007). Its plasticity manifests bidirectionally in response to a separate set of bidirectional modifications in synaptic plasticity (Belmeguenai et al., 2010; Shim et al., 2017; Yang & Santamaria, 2016). This is indicative of homeostatic maintenance within the neural circuitry (Turrigiano, 2012) and supports the notion that multiple mechanisms operate within the cerebellar memory system (Boyden et al., 2004). Notably, OKR learning and eye-blink conditioning are accompanied by the down-regulation and upregulation of PC intrinsic excitability (PC-IE), respectively (Kim & Kim, 2020; Titley et al., 2020). Transgenic mouse models with intact synaptic transmission but deficient intrinsic plasticity show impaired consolidation despite successful learning acquisition (Jang et al., 2020; Ryu et al., 2017). In different regions such as the hippocampus, increased cortical excitability has been observed in response to eye-blink and fear conditioning (Kuo et al., 2008; Mckay et al., 2009). Although numerous studies have established a correlation between the level of intrinsic excitability and the quality of learning and memory, research exploring the direct impact of neuronal IE on the formation of long-term memory remains limited. This knowledge gap is particularly pronounced in the cerebellum, as the potential implications of manipulating PC-IE on memory consolidation remain unknown. Evidence shows that PC-IE fluctuates in a time-dependent manner following motor learning (Jang et al., 2020), but the temporal dependency of its contribution to consolidation remains unclear. Therefore, it is imperative to establish causality between consolidation and PC-IE and to determine the optimal timing of PC-IE for effective long-term storage.

Inter-regional communication between the cerebellar cortex and the nuclei underlies systems consolidation (Kassardjian et al., 2005; Shutoh et al., 2006), and it has been suggested to be accompanied by PC-dependent synaptic plasticity and intrinsic excitability of neurons in the MVN (Jang et al., 2020; McElvain et al., 2010). Given this, cortical PC is a plausible neural substrate for connecting two different brain regions. However, the direct impact of changes in PC-IE on systems consolidation regarding the intrinsic excitability of FTNs remains elusive.

In the present study, we hypothesized that PC-IE is essential for the maturation of motor memories and the formation of neural links between the cortex and the nuclei during the consolidation period. Using neuron-specific optogenetic manipulations in conjunction with the OKR paradigm, we provide compelling evidence that PC-IE plays a crucial determinant in facilitating memory consolidation, as we observed that upregulation of cortical PC-IE within a specific time window results in impaired memory consolidation. Moreover, we demonstrate that PC-IE manipulation compromises the intrinsic plasticity of subsequent FTNs in the nucleus, suggesting that the PC-FTN circuit is highly dependent on PC-IE. Our findings provide a novel perspective on the non-synaptic mechanism underlying the systems consolidation process in the cerebellar circuit, which goes beyond the traditional perspective of synaptic plasticity.

## Results

### Optogenetic manipulation of PC-IE increases spontaneous firing

Recent studies have demonstrated that the intrinsic excitability of PCs is strongly associated with cerebellar motor learning (Jang et al., 2020; Kim & Kim, 2020; Titley et al., 2020), but yet, the actual causality between PC-IE and motor memory consolidation through cell-type specific manipulation remains unexplored. Hence, we devised an appropriate method for the selective manipulation of PC-IE in the flocculus. To achieve this, we employed optogenetics using the stabilized step-function opsin (SSFO) (Yizhar et al., 2011). It promptly elicits depolarization and stable excitatory state in neurons in response to optostimulation. The experimental design involved the bilateral injection of AAV1-Ef1a-DIO hChR2(C128S/D156A)-EYFP or AAV1-Ef1a-DIO EYFP into the flocculi of Pcp2-cre (L7-Cre) mice, which allowed specific recombinase activity in the floccular PC layer (Figure 1a). Following the surgery, an in-vitro whole-cell patch clamp recording of PCs was conducted (Figure 1b). We validated that in response to 473 nm optostimulation, which activates the opsin, the spontaneous firing of SSFO-expressing PCs was significantly elevated (Figure 1c). Specifically, the average firing rate of SSFO-expressing PCs was robustly enhanced, whereas EYFP-expressing PCs did not show any changes (Figure 1d). Furthermore, upon 593 nm optostimulation, which deactivates the opsin, the elevated firing rate of SSFO-expressing PCs reverted to the basal condition, and no change was detected in EYFP-expressing PCs, as expected (Figure 1c, d). Moreover, our ex-vivo result indicated that PCs of optostimulated SSFO-expressing mice following motor learning exhibited significantly higher IE than that of EYFP-expressing mice (Supplementary Figure 1a, b). These results establish the efficacy of the proposed approach for selectively manipulating PC-IE during the consolidation period, providing a promising foundation for investigating the causal relationship between PC-IE and cerebellar memory consolidation.

**Figure 1.**
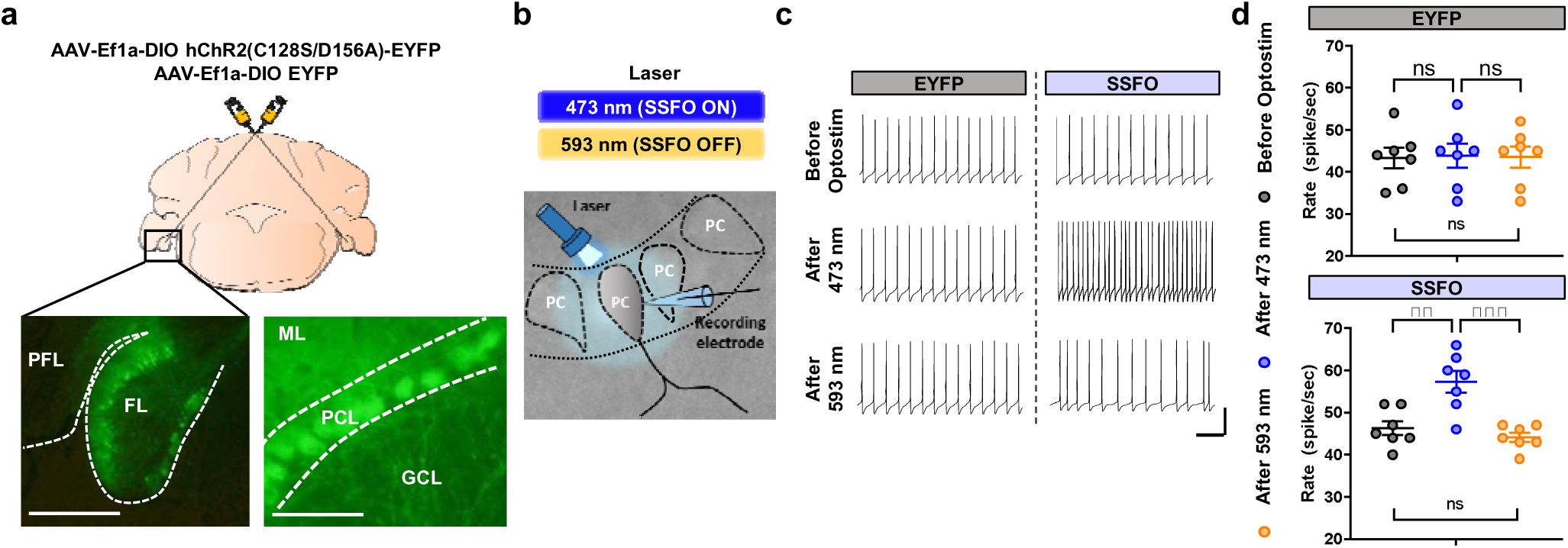
Optogenetic manipulation of PC-IE increases spontaneous firing. **a.** Schematic diagram of the virus injection for the optogenetic manipulation of floccular Purkinje cells (PCs). Bottom left, the virus is specifically expressed in PCs of the flocculus (FL), which is located below the paraflocculus (PFL) (scale bar indicates 500 µm). Bottom right, the granule cell layer (GCL), PC layer (PCL), and molecular layer (ML) are shown (scale bar indicates 150 µm). **b.** Scheme for optogenetic activation/deactivation of the SSFO virus expressed in PCs using a blue laser (473 nm)/yellow laser (593 nm). **c.** Representative traces of spontaneous PC firing. Traces of both the EYFP and SSFO groups are shown under three different conditions: before optostimulation, after 473 nm optostimulation, and after 593 nm optostimulation (vertical scale bar indicates 20 mV and horizontal scale bar represents 100 ms). **d.** Spontaneous firing rate of SSFO-expressing PCs (n=7) and EYFP-expressing PCs (n=7). The rate of SSFO-expressing PCs significantly increased and decreased following 473 nm and 593 nm optostimulation, respectively (SSFO, ** p = 0.0017 and *** p = 0.0003; one-way ANOVA Tukey’s multiple comparisons). The error bars indicate ±SEM.

### Optogenetic manipulation of PC-IE impairs memory consolidation

To understand the contribution of intrinsic excitability to memory consolidation, we employed optogenetics during the behavioral assessment. Specifically, we utilized the OKR protocol to train mice in adapting their ocular reflex in response to visual stimulation. To manipulate the floccular PC excitability *in vivo* during the consolidation period, we bilaterally implanted an optic cannula into the flocculi of mice that had undergone surgery for SSFO/EYFP expression. One mouse underwent 2 days of training. On the first day, it was trained for 50 min and placed back into the home-cage in a completely dark condition to rest without any visual interruption (Figure 2a, top panel). During the training session, an animal was subjected to continuous oscillation (±5° peak-to-peak motion at 0.5 Hz) of the optokinetic screen designed by vertical stripe patterns (Figure 2b). Once training was completed, bilateral 473 nm optostimulation was administered at 0 min post-learning to activate the SSFO and another set of bilateral 593 nm optostimulation was provided at 24 hr post-learning to deactivate the SSFO (Figure 2a, bottom panel). Proper training reduced the retinal slip, which directly reflected the accomplishment of motor learning in both the SSFO and EYFP groups (Figure 2c, pre- and post-learning). When mice were exposed to continuous sinusoidal screen oscillation before training, the gain, a value representing how closely the eye motion overlaps with the rotation of a screen, ranged between 0.22 and 0.58 (EYFP = 0.235-0.576; SSFO = 0.217-0.557); after training, the gains were increased ranging between 0.38 and 0.84 (EYFP = 0.571-0.746; SSFO = 0.381-0.838) (Figure 2d). Improved oculomotor performance was observed in both groups. After confirming the learning results, we activated the SSFO during the initial phase of the consolidation period. Thus, we were able to instantly disrupt the learning-induced depression of intrinsic excitability that is reported to be promptly exhibited after cerebellar motor learning (Jang et al., 2020; Kim & Kim, 2020). As a result, long-term memory was normally formed in the EYFP group after 24 hr of learning, but was abolished in the SSFO group due to aberrant conditions in the neural circuitry for consolidation (Figure 2d). Specifically, the consolidation percentage of motor memory in the SSFO group was significantly lower than that in control group (Figure 2e). To investigate the potential disruption of memory consolidation attributed to the mere expression of SSFO, we conducted an additional set of OKR learning using mice expressing SSFO. Subsequently, these mice, which had previously showed impaired consolidation with optogenetic manipulation, underwent for a recovery period of 2 to 3 weeks. Mice with relatively high gains before the second set of learning, indicative of the remaining effects from the first learning set, were excluded from the second assessment. When SSFO-expressing mice were trained and subjected to a consolidation period without interference from optogenetic manipulation, their long-term memory was steadily retained (Supplementary Figure 2a, b). Therefore, these results reveal the importance of the proper expression of PC-IE in facilitating memory consolidation.

**Figure 2.**
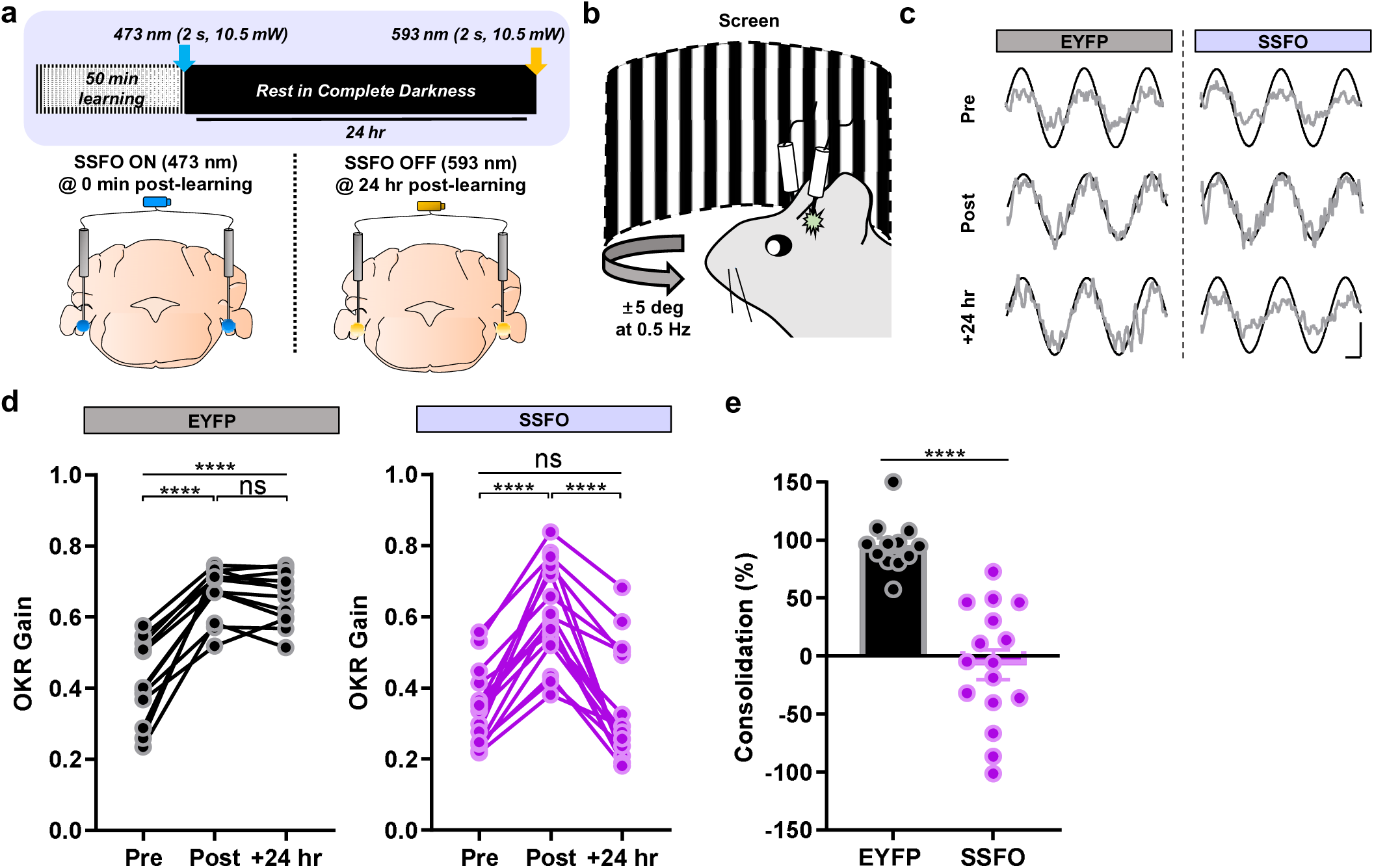
Optogenetic manipulation of PC-IE impairs memory consolidation. **a.** Top, experimental scheme for the optokinetic reflex (OKR). On the first day after learning was complete, 473 nm optostimulation (2 s, 10.5mW) was administered to the mouse via an optic cannula at 0 min of the post-learning period. The mouse was then placed in a cage to rest in complete darkness for 24 hr. On the second day, 593 nm optostimulation (2s, 10.5 mW) was administered to the mouse, and the remaining gain was measured. Bottom, schematic diagram shows the location of optic cannula implantation, which is the flocculus of the cerebellum, and the settings of optostimulation. **b.** Illustration of the OKR behavioral test. During the learning period, visual stimulation was provided by the horizontal rotation of a screen at a constant speed (±5 deg at 0.5 Hz) and the table was stationary. **c.** Representative traces of the screen (black sinusoidal curve) and eye movements (grey sinusoidal curve). Traces of both the EYFP and SSFO groups were presented at three different periods: pre (before learning), post (after learning) and +24 hr (post 24 hr). The vertical scale bar represents 10 degrees per second and horizontal scale bar represents 0.5 second. **d.** Gain changes in the EYFP and SSFO groups following learning. The gain values in both groups (EYFP, n = 14 mice; SSFO, n = 16 mice) significantly increased after learning (EYFP, **** p < 0.0001; SSFO, **** p < 0.0001; repeated-measure [RM] one-way analysis of variance [ANOVA], Tukey’s multiple comparisons). After 24 hr of consolidation period, the EYFP group showed the robust maintenance of long-term memory (p = 0.4122; RM one-way ANOVA Tukey’s multiple comparisons), whereas the SSFO group shows impaired consolidation (**** p < 0.0001; RM one-way ANOVA, Tukey’s multiple comparisons). **e.** Comparison of the consolidation percentages between the EYFP and SSFO groups. The consolidation percentage in the EYFP group is significantly higher than that in the SSFO group (EYFP, n = 14 mice; SSFO, n = 16 mice; Unpaired t test, **** p < 0.0001). The error bars indicate ±SEM.

### PC-IE contributes to the formation of long-term memory during the early phase of the consolidation period

In addition to OKR learning, long-term memories of other cerebellum-dependent behaviors, such as the vestibulo-ocular reflex (VOR) and eye-blink conditioning are susceptible to interference within specific periods of consolidation. When the intracortical activity was disrupted between 45 and 90 min after eye-blink conditioning, the rabbits were unable to properly express retained long-term memory (Cooke et al., 2004). Similar to OKR, VOR requires the cerebellum for visual stabilization when stimulated by vestibular inputs. After the cats were trained for the VOR, they were subsequently exposed to a condition in which their vision was being constantly interrupted during the first hour of the consolidation period. Following the visual interruption, the cats exhibited reverted learning outcomes (Titley et al., 2007). In addition, consolidation was prevented when G-protein coupled receptors that are involved in learning and memory were pharmaceutically suppressed within 1hr after the learning period (Steinmetz & Freeman, 2016).

Building upon the findings from previous studies, the primary objective of this study was to ascertain whether a critical time window exists during which PC-IE plays a pivotal and highly effective role in memory consolidation. Given our observation that optogenetic manipulation throughout the entire consolidation period resulted in the loss of long-term memory, we implemented two distinct behavioral schemes involving different time points of optogenetic manipulation. Some mice were given 473 nm optostimulation at 0 min post-learning (Figure 3a, *Scheme #1*), and a break of approximately 2 to 3 weeks was provided to ensure the extinction of previously acquired motor memory for a revised scheme of the next experiment. Subsequently, they underwent a second trial of learning, with optostimulation at 90 min post-learning (Figure 3a, *Scheme #2*). The order of manipulation was counterbalanced to eliminate any potential bias. Consequently, regardless of the behavioral scheme, the EYFP group demonstrated robustly sustained long-term memory, as shown by the high consolidation percentage, in both tests (Figure 3b, c). However, the SSFO group exhibited varying outcomes based on the timing of the manipulation. When SSFO was activated to increase PC-IE at 0 min post-learning, the previously enhanced gain levels were reversed, resulting in a relatively low consolidation percentage (Figures 3d [left] and 3e). By contrast, when the manipulation was administered 90 min after learning, the increased gain was maintained, resulting in a high consolidation percentage (Figures 3d [right] and 3e). Outside this critical time window, alterations in PC-IE did not significantly affect formation of long-term memory, emphasizing the pivotal role of the timing of PC-IE contribution to memory consolidation. Furthermore, we substantiated the significance of 90 min post-learning period by optogenetically manipulating PC-IE exclusively within this initial timeframe. Despite any interference, the EYFP group exhibited intact memory consolidation (Figures 3f [left] and 3g), but, as anticipated, the SSFO group was incapable of retaining long-term memory (Figure 3f [right] and 3g), highlighting that the induction of aberrant PC-IE was powerful enough to disturb the nature of learning and memory. These compelling results clearly demonstrate that PC-IE exercises physiological leverage to facilitate memory consolidation within 90 min post-learning period.

**Figure 3.**
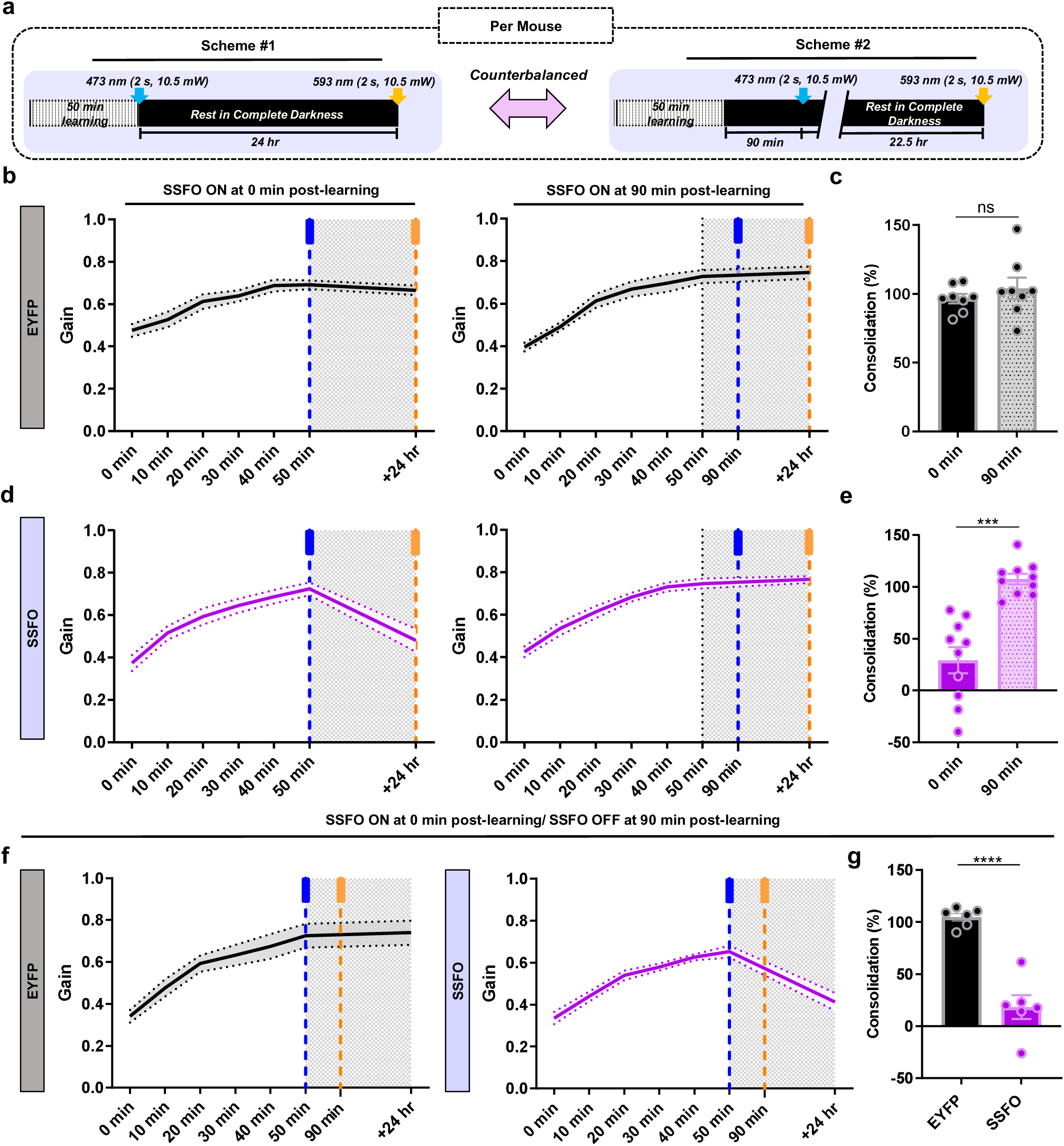
PC-IE contributes to the formation of long-term memory during the early phase of the consolidation period. **a.** Two different experimental schemes for OKR. *Scheme #1*, 50 min of learning was conducted on the first day; once the learning was complete, 473 nm optostimulation (2 s, 10.5mW) was administered to mouse via an optic cannula at 0 min post-learning period. The mouse was then placed back in a cage to rest in complete darkness for 24 hr. On the second day, 593 nm optostimulation (2s, 10.5 mW) was given to a mouse, and remaining gain was measured. *Scheme #2*, same behavior protocol for OKR was conducted as *scheme #1*, but 473 nm optostimulation (2 s, 10.5mW) was given at 90 min post-learning period. **b.** OKR learning curves from 0 to 50 min and +24 hr of the EYFP group under two different conditions. Left, 473 nm optostimulation (2 s, 10.5mW) was administered to the mouse via an optic cannula at 0 min post-learning period. Right, the same treatment was administered at 90 min post-learning period. **c.** Comparison of consolidation percentages of EYFP-expressing mice under two different conditions. The consolidation percentages of EYFP-expressing mice optostimulated at 0 min post-learning period are comparable to those of EYFP-expressing mice optostimulated at 90 min post-learning period (0 min, n = 8; 90 min, n = 8; Paired t test, p = 0.3248) **d.** OKR learning curves from 0 to 50 min, and +24 hr of the SSFO group under two different conditions. Left, 473 nm optostimulation (2 s, 10.5mW) administered a mouse via an optic cannula at 0 min post-learning period. Right, the same treatment was administered at 90 min post-learning period. **e.** Comparison of consolidation percentages of SSFO-expressing mice under two different conditions. The consolidation percentages of SSFO-expressing mice optostimulated at 90 min post-learning period are significantly higher than those of SSFO-expressing mice optostimulated at 0 min post-learning period (0 min, n = 10; 90 min, n = 10; paired t-test, *** p < 0.0010). **f.** OKR learning curves from 0 to 50 min and +24 hr. Left, learning curve of the EYFP group: 473 nm optostimulation (2 s, 10.5mW) was administered to the mouse via an optic cannula at 0 min post-learning period, and 593 nm optostimulation (2s, 10.5 mW) was administered at 90 min post-learning period. Right, learning curve of the SSFO group: 473 nm optostimulation (2 s, 10.5mW) was administered to the mouse via an optic cannula at 0 min post-learning period, and 593 nm optostimulation (2 s, 10.5mW) was administered at 90 min post-learning period. **g.** Comparison of consolidation percentages of between the EYFP (n = 6) and SSFO (n = 6) groups. The consolidation percentages of the SSFO group was significantly lower than that of the EYFP group (unpaired t-test, **** p < 0.0010). The error bars indicate ±SEM.

### Dynamics of PC-IE within the critical temporal window following motor learning

Under general circumstances, PC-IE remains constant, but according to a previous study, VOR learning induces the temporal adjustment of PC-IE throughout the consolidation period in a bidirectional manner, which consists of decreased or restored levels of excitability depending on the timeline (Jang et al., 2020; Kim & Kim, 2020). Kim et al. (2020) found that OKR learning reduces PC-IE immediately after the completion of motor learning (Kim & Kim, 2020). However, the fluctuation of excitability at various time points has not been thoroughly elaborated. Therefore, it remains unclear how PC-IE dynamics are exhibited after OKR learning, particularly within the observed critical time window. To address this question, we investigated the underlying cellular mechanisms that establish a critical time window. Once the mice were fully trained for OKR, we prepared brain slices to perform whole-cell patch clamp recordings from floccular PCs at various post-learning periods, which allowing us to explore the dynamics of PC-IE during consolidation (Figure 4a). Some slices were acquired at 0 min post-learning, while others were prepared at 90 min post-learning. To evaluate PC-IE, we provided current injections from +0 pA to +900 pA (100 pA increment per injection, 500 ms long) in the presence of NBQX (2,3-dihydroxy-6-nitro-7-sulfamoyl-benzo[*f*]quinoxaline) and picrotoxin to block excitatory and inhibitory synaptic inputs, respectively. The results demonstrated that the mean firing rate of floccular PCs was significantly reduced compared with that of naïve control conditions, but after 90 min post-learning, the rate recovered to that of the naïve condition (Figure 4b, c). This indicates that the depression of PC-IE did not last for more than 90 min.

**Figure 4.**
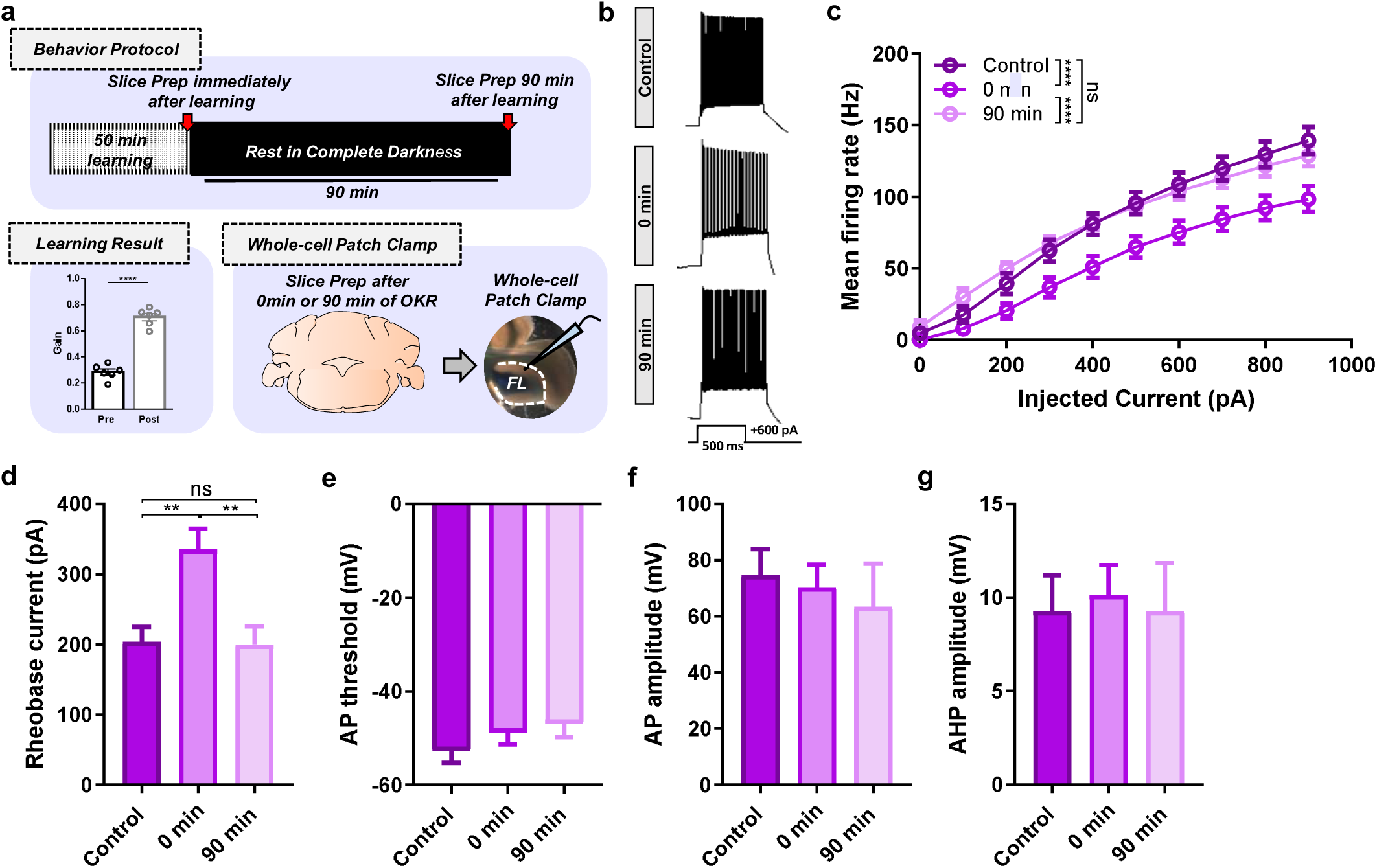
Dynamics of PC-IE within the critical temporal window following motor learning. **a.** Top, the ex-vivo experimental scheme for OKR and whole-cell patch clamp recording. After 50 min of learning, slice preparation for whole-cell patch clamp recording was conducted at 0 min post-learning and 90 min post-learning periods. Bottom left, learning-induced increase in the OKR gain is presented (mouse, n = 5; paired t -test, *** p = 0.004). Bottom right, coronal slices are acquired, and electrophysiological recordings are conducted in the flocculus. **b.** Raw traces of PC firing under 600 pA current injection under three different conditions: control, at 0 min post-learning period, and at 90 min post-learning period. **c.** The mean firing rate (Hz) is significantly decreased in the learned group at 0 min post-learning compared with that in the control (naïve) and 90 min post-learning groups (control, n = 14; 0 min, n = 14; 90 min, n = 10; two-way analysis of variance [ANOVA] with Tukey’s multiple comparisons, **** p < 0.0001). **d.** OKR learning elicits significant increase in the rheobase current at 0 min post-learning period (One-way ANOVA multiple comparisons, 0 min vs. 90 min, * p = 0.0336; 0 min vs. Control, * p = 0.0187). **e.** No significant differences in the AP threshold. **f.** No significant differences in the AP amplitude. **g.** No significant differences in the AHP amplitude. The error bars indicate ±SEM.

Furthermore, we analyzed the changes in action potential (AP) properties to explain the depression of PC-IE. We measured four AP parameters: the rheobase current, AP threshold, AP amplitude and afterhyperpolarization (AHP) amplitude. Among these parameters, only the rheobase current exhibited significant changes. It was significantly increased at 0 min post-learning, but lasted temporarily such that the increased current was reduced back to the basal level at 90 min post-learning (Figures 4b, c). This implies that, within 90 min post-learning period PC-IE decreased and required a larger current threshold to fire an AP. Beside the rheobase current, other properties were not altered following OKR learning (Figure 4d-f).

To assess whether our optogenetic manipulation at 0 min post-learning induced an abnormal condition, thereby blocking the learning-induced depression in PC-IE. The ex-vivo results revealed that PCs of optostimulated SSFO-expressing mice, following motor learning, exhibited significantly higher firing rates than those of EYFP-expressing mice (Supplementary Figure 1a, b). This heightened activity directly signifies the abolished depression in PC-IE of the SSFO group. Consequently, the loss of long-term memory can be attributed to the prevention of learning-induced depression in PC-IE, a process anticipated to occur after learning.

### Disrupted plasticity of intrinsic excitability of flocculus-targeting neurons in the MVN

Postsynaptic circuitry of cortical PCs is primarily organized by neurons in the MVN, characterized by two distinct types: flocculus-targeting neurons (FTNs) and nonFTNs (Bagnall et al., 2007; Sekirnjak & du Lac, 2006). Among these types, FTNs, specifically, play an active role in facilitating memory consolidation (Kassardjian et al., 2005; Matsuno et al., 2016; Shin et al., 2011; Shutoh et al., 2006), as they serve as the subsequent destinations for memory transfer. The ability of FTNs to maintain and express long-term memory is supplemented by both synaptic and intrinsic plasticities (Gittis & du Lac, 2006; McElvain et al., 2010). Previous studies reported that not only intrinsic plasticity of PC was elicited following motor learning, but also a potentiation of intrinsic excitability was induced in VN neurons (Jang et al., 2020). Therefore, we investigated whether optogenetic manipulation of PC-IE directly influences the expression of intrinsic plasticity of FTNs. Pcp2-cre (L7-Cre) mice underwent surgery for SSFO and EYFP injections into the flocculus. These mice underwent OKR learning, followed by optostimulation to activate SSFO for the entire period of consolidation, and eventually, acute slices were obtained at 24 hr post-learning period for whole-cell patch clamp recording (Figure 5a, top). Rich expression of SSFO/EYFP in floccular PCs allowed for the morphological distinction of FTNs and nonFTNs, indicated by axon terminals with multiple boutons en passant densely surrounding neurons in MVN (Figures 5a [bottom] and 5b).

**Figure 5.**
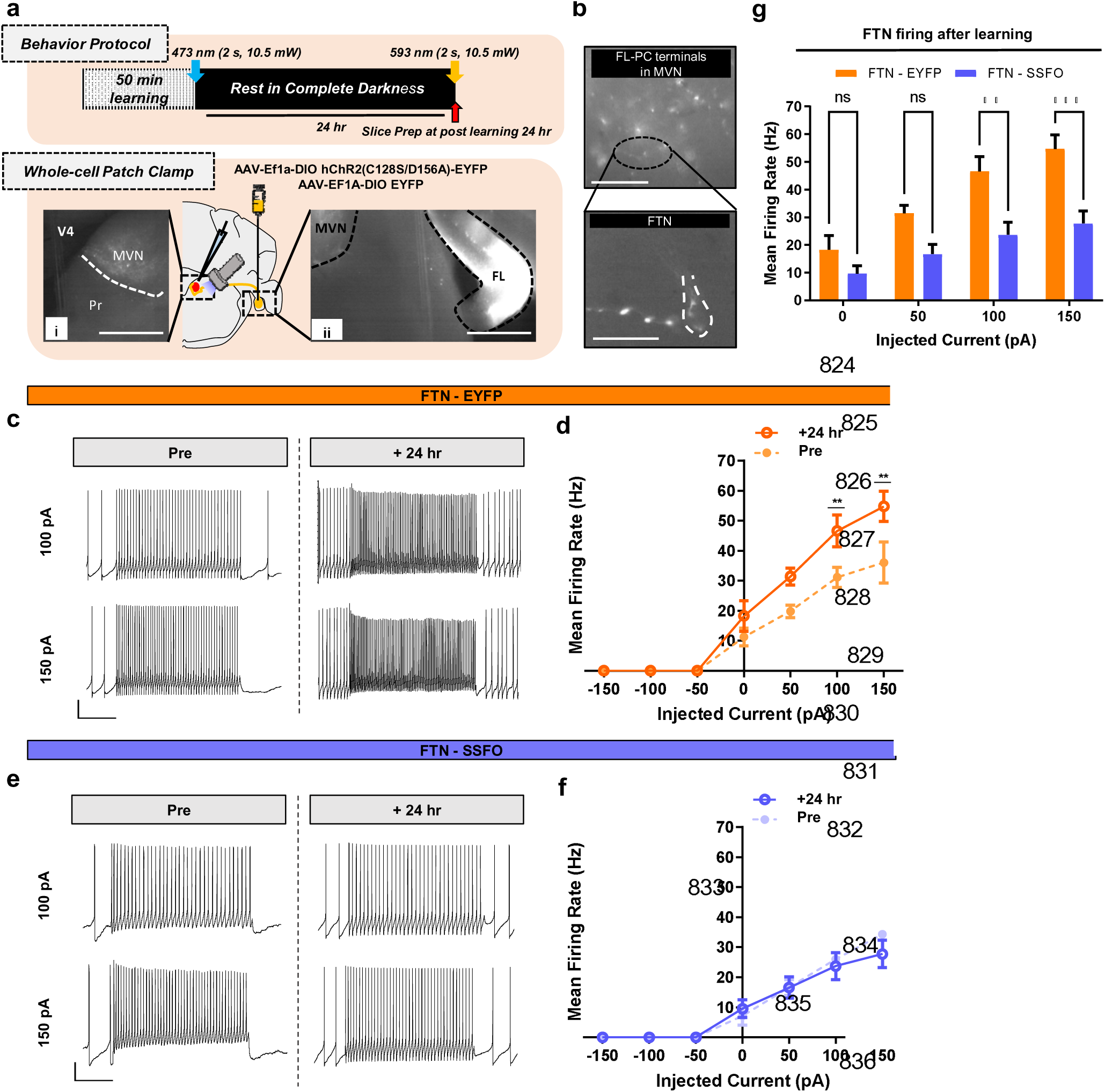
Disrupted plasticity of intrinsic excitability of flocculus-targeting neurons in the MVN. **a.** Top, an ex-vivo experimental scheme for OKR and whole-cell patch clamp recording. After 50 min of learning, slice preparation for whole-cell patch clamp recording was conducted at 24 hr post-learning period. Bottom (i), coronal slices are acquired, and electrophysiological signals of FTNs in the medial vestibular nucleus (MVN) were recorded (scale bars indicate 500 µm). (ii), The virus expression (EYFP/SSFO) in the flocculus is validated (scale bars indicate 500 µm). **b.** Top, an image of floccular PC terminals expressed in the MVN (scale bar indicates 25 µm). Bottom, an image of FTN with *boutons en passant* (scale bar indicates 20 µm). **c.** Raw traces representing the excitability of EYFP-expressing FTNs at 100 pA and 150 pA current injections under learned (n = 6) and naïve (n = 7) conditions. The vertical scale bar represents 10 mV and the horizontal scale bar represents 200 ms. **d.** Comparison of intrinsic excitability of EYFP-expressing FTNs under naïve and learned conditions. Excitability at 100 pA and 150 pA current injections is significantly higher in the learned group (n = 6) than in the naïve group (n = 7) (Two-way analysis of variance [ANOVA] with Sidak’s multiple comparisons: at 100 pA, ** p = 0.0065; at 150 pA, ** p = 0.0025). **e.** Raw traces representing the excitability of SSFO-expressing FTNs at 100 pA and 150 pA current injections under learned (n = 7) and naïve (n = 6) conditions. The vertical scale bar represents 10 mV and the horizontal scale bar represents 200 ms. **f.** Comparison of intrinsic excitability of SSFO-expressing FTNs under naïve and learned conditions. The excitability of the learned (n = 7) and naïve (n = 6) groups is (n = 6) comparable. They exhibit no significant difference. **g.** Comparisons of mean firing rates between the FTNs of the EYFP and SSFO groups after learning. EYFP-expressing FTNs (n = 6) exhibit significantly higher mean firing rates than SSFO-expressing FTNs (n = 7) at 100 pA and 150 pA current injections (two-way ANOVA with Sidak’s multiple comparisons: at 100 pA, ** p = 0.0020; at 150 pA, *** p = 0.0002). The error bars indicate ±SEM.

To further investigate whether learning-induced plasticity of intrinsic excitability occurs in the FTNs, we measured the mean firing rates of the EYFP and SSFO groups under both naïve and learned conditions. Comparisons revealed distinct patterns. In EYFP-expressing mice, intact memory consolidation post-learning was accompanied by the potentiation of intrinsic excitability in *in-vitro* (Figures 5c, d). Conversely, in SSFO-expressing mice, the abnormal increase in PC-IE resulted in impaired memory consolidation and the absence of intrinsic plasticity in FTNs (Figures 5e, f). Upon completion of the learning and subsequent consolidation period, significant disparities in mean firing rates between EYFP-expressing and SSFO-expressing FTNs were particularly evident during 100 pA and 150 pA stimulations (Figure 5g).

Turning our attention toward nonFTN circuitry, which remains isolated from the cortical learning circuit, we speculated that there would be no discernible learning-induced plasticity in nonFTNs following motor learning. As anticipated, motor learning did not elicit intrinsic plasticity in nonFTNs in either the EYFP or SSFO groups, as there was no significant difference in the mean firing rates (Supplementary Figures 3a-d). Unlike FTNs, nonFTNs in both the EYFP and SSFO groups showed comparable excitability following motor learning (Supplementary Figure 3e). Overall, these data clearly demonstrate that the proper maintenance of long-term memory involves the expression of intrinsic plasticities in both PCs and FTNs, forming the foundation for systems consolidation, which, in turn requires inter-regional communication.

## Discussion

Conventional theories of learning and memory have predominantly focused on bidirectional changes in synaptic efficacy at multiple sites within the cerebellar circuitry(Boyden et al., 2004). However, our findings challenge this view by revealing a crucial role for IE in PCs during systems memory consolidation. IE reflects a neuron’s capacity to generate action potentials independently of synaptic input (Chen et al., 2020), and its role in memory consolidation has remained largely unexplored. To address this knowledge gap, we employed a comprehensive approach that integrates optogenetics, behavioral assays, and electrophysiological recordings. Optogenetic manipulation of PC-IE resulted in an aberrant increase in neuronal excitability and spontaneous firing, resulting in a discernible impairment in the consolidation of motor memory. These findings redefine our understanding of cerebellar memory circuitry, emphasizing that memory consolidation is not solely dependent on synaptic plasticity but also on the precise regulation of intrinsic neuronal properties. Notably, cerebellar learning induces time-dependent fluctuations in PC-IE, particularly following VOR. Contrary to previous reports that highlighted a transient decrease in PC-IE shortly after OKR (Jang et al., 2020; Kim & Kim, 2020), our detailed temporal analysis uncovers sustained changes in excitability that persist well beyond the initial learning phase. A pronounced decrease in PC-IE underscores an important temporal window that decisively influences the achievement of memory consolidation. Importantly, these changes in IE are not confined to the cerebellar cortex but extend to downstream targets, such as FTNs in the MVN. Induction of intrinsic plasticity in the nuclei was prevented due to the abnormal expression of PC-IE. This suggests that the interplay between cortical and nuclear region, mediated by enduring changes in IE, is essential for systems consolidation. This concept aligns with the hypothesis that intrinsic plasticity may perform as a trigger or conductor of systems-level consolidation within the circuits involving the medial prefrontal cortex-basolateral amygdala, and hippocampus (Lisman et al., 2018; Song et al., 2015).

The involvement of intrinsic neuronal excitability in memory consolidation has been often observed in other memory systems (Daoudal & Debanne, 2003; Kuo et al., 2008; Mckay et al., 2009; Zhang & Linden, 2003). Interestingly, our data revealed that cerebellar learning elicited depression in PC-IE, whereas neurons in the amygdala exhibited increased neuronal excitability associated with fear learning (Zhou et al., 2009). This divergence underscores the unique mechanism governing cerebellar memory consolidation and suggests that the cerebellum employs distinct strategies for different types of learning and memory. Hence, optogenetic manipulation of exciting the PCs in this study may have disturbed the incomparable nature of the cerebellum.

A particular novel aspect of our work is the identification of 90-minute critical window following motor learning, during which PC-IE is dynamically regulated. This temporal constraint suggests that the initial phase of consolidation is a critical determinant for transforming short-term memory into long-term memory. Our electrophysiological recordings demonstrated that PC-IE at 0 min post-learning period was significantly decreased compared with that of the naïve condition, which gradually returns to the baseline within this critical period. Given the role of PC firing as the primary output of the cortex, encoding temporal and sensory information (De Zeeuw et al., 2011), we propose that fluctuations in PC-IE is a plausible mechanism for determining the critical time window that is integral to the consolidation process. This finding parallels the early phase of hippocampal systems consolidation, where synaptic plasticity facilitates the formation of long-term memory at earlier period of consolidation (Goto et al., 2021). Moreover, although transient fluctuations of hippocampal excitability have been suggested to conduct memory consolidation (Thompson et al., 1996), this study represents the first detailed exploration of the critical time window in cerebellum-dependent memory consolidation concerning the dynamics of neuronal intrinsic excitability.

It is widely recognized that memory consolidation requires de novo protein synthesis, and the temporal aspect of protein synthesis is of paramount importance for long-term storage (Shafik et al., 2022; Shrestha & Klann, 2022). Long-term contextual fear memory formation in the hippocampus, for instance, relies on transcription factors and associated signaling cascades, such as the cyclic AMP response element binding protein and the Ras/Erk pathway, both of which are essential for protein synthesis (Athos et al., 2002; Tian et al., 2004; Wood et al., 2005). Intriguingly, cerebellum-dependent motor learning is also associated with certain modifications in protein expression, though the specific proteins involved differ depending on whether they are linked to short- or long-term memory (Kim et al., 2020). Our data emphasize the existence of a critical time window for memory consolidation regulated by intrinsic excitability. Nonetheless, we acknowledge that we have not yet provided further insights into the molecular mechanisms underlying this phenomenon. As such, the involvement of protein synthesis in establishing this critical time window and its relationship with fluctuations in PC-IE are areas that require further investigation.

Similar to the systems consolidation processes observed in the hippocampus, where remote memories are established in the neocortex, the cerebellar cortex-to-nuclei network also undergoes analogous mechanisms. Previous studies suggest that hippocampal and neocortical synaptic plasticity share conserved molecular substrates and conditioned pathways (Kirkwood et al., 1993). Moreover, hippocampal modifications, such as conformational changes in dendritic spines and suppression of neuronal output, directly influence cortical representations (Restivo et al., 2009; Tanaka et al., 2014; Vetere et al., 2011), but unlike studies on the hippocampus, there is limited understanding of the neural properties underlying cerebellar systems consolidation. Therefore, we further investigated the post-circuitry of floccular PCs, which primarily interface with FTNs in the MVN. Our findings reveal that PC excitability and the intrinsic excitability of FTNs in the MVN play an important role in the construction of cerebellum-dependent long-term memory. Disruption of PC-IE was also accompanied by the impaired plasticity of the intrinsic excitability of FTNs, indicating a direct coupling of intrinsic plasticities across distinct regions within a unified memory circuit. Our data show that the intrinsic plasticity of PCs provides guidance to drive the induction of intrinsic plasticity in FTNs, which aligns with previous models proposing that PC activity modulates IE of FTNs (Jang et al., 2020; McElvain et al., 2010). Additionally, we extended our analysis to another prominent group of neurons in the MVN, nonFTNs. Given that nonFTNs receive direct inputs from vestibular organs and lack synaptic connections with floccular PCs, it was unsurprising that alterations in PC-IE did not affect the IE of nonFTNs

PCs, are a rare and exclusive type of neuron found solely within the cerebellar cortex, and have long been associated with motor coordination and movement (Hoogland et al., 2015; Thach, 1998). Overall, this study demonstrated that the ability to fine-tune firing patterns in a time-dependent manner by modulating intrinsic excitability plays an essential role in memory consolidation. This mechanism is crucial for the precise refinement of motor memory, prompting us to explore the fundamental question of why PCs employ such sophisticated approaches. The cerebellum’s capacity to process vast amounts of sensory input through a single output pathway, the PCs, raises intriguing questions about the evolutionary and functional significant of IE in this context (Voogd & Glickstein, 1998). Despite the abundance of sensory inputs, only one pathway of cerebellar cortical output exists: the PCs, which exercise a far-reaching influence on numerous regions in the cerebral cortex to modify both motor and non-motor behaviors(Carta et al., 2019; Li et al., 2023; Low et al., 2021). Although PCs in different cerebellar lobules exhibit no morphological differences, compelling evidence demonstrates that each lobule serves specific roles by encoding different sensory modalities (Barmack et al., 1992; Barmack et al., 1993; Kim et al., 2009); hence, PCs exhibit different firing patterns with distinct membrane properties according to their location (Kim et al., 2012). In addition to the results of previous studies, the results of this study led us to speculate that PCs may have evolved to establish specific patterns of spike activity to conduct a diverse range of cerebellum-dependent tasks, thereby optimizing the biological system and allowing for efficient performance. Our findings suggest that these specialized firing patterns are, in part, shaped by the dynamic regulation of IE, which optimizes cerebellar output for the consolidation of motor memories.

The construction of computational models of neuronal pathways has proven to be a useful tool for analyzing and understanding memory circuits (Rizza et al., 2021; Yamazaki & Nagao, 2012; Yamazaki et al., 2015). Based on our findings, future studies may apply these insights to computational modeling and simulations to predict the patterns of PC activity during different behavior assessments across multiple time points. This approach has the potential to provide deeper insights into the intricate interplay between biological and computational mechanisms governing the precise tuning of PC firing through intrinsic excitability. Moreover, extending the application of these computational models to broader memory systems across diverse brain regions offers an opportunity to expand our understanding of memory consolidation processes beyond the cerebellum and a unified understanding of memory as a fundamental cognitive process.

In summary, our investigation provides a paradigm shift in the understanding of cerebellar memory consolidation, highlighting the critical role of PC-IE, particularly within the critical 90-minute time window. We demonstrate that memory consolidation within the cerebellum is not solely a function of synaptic plasticity, but also requires precise temporal regulation of neuronal excitability. These findings have broad implications for the study of memory across the brain and offer potential therapeutic targets for disorders involving cerebellar dysfunction (Bruchhage et al., 2018; Moreno-Rius, 2019; Shakiba, 2014; Wu & Hallett, 2013).

## Materials and methods

### Key resources table

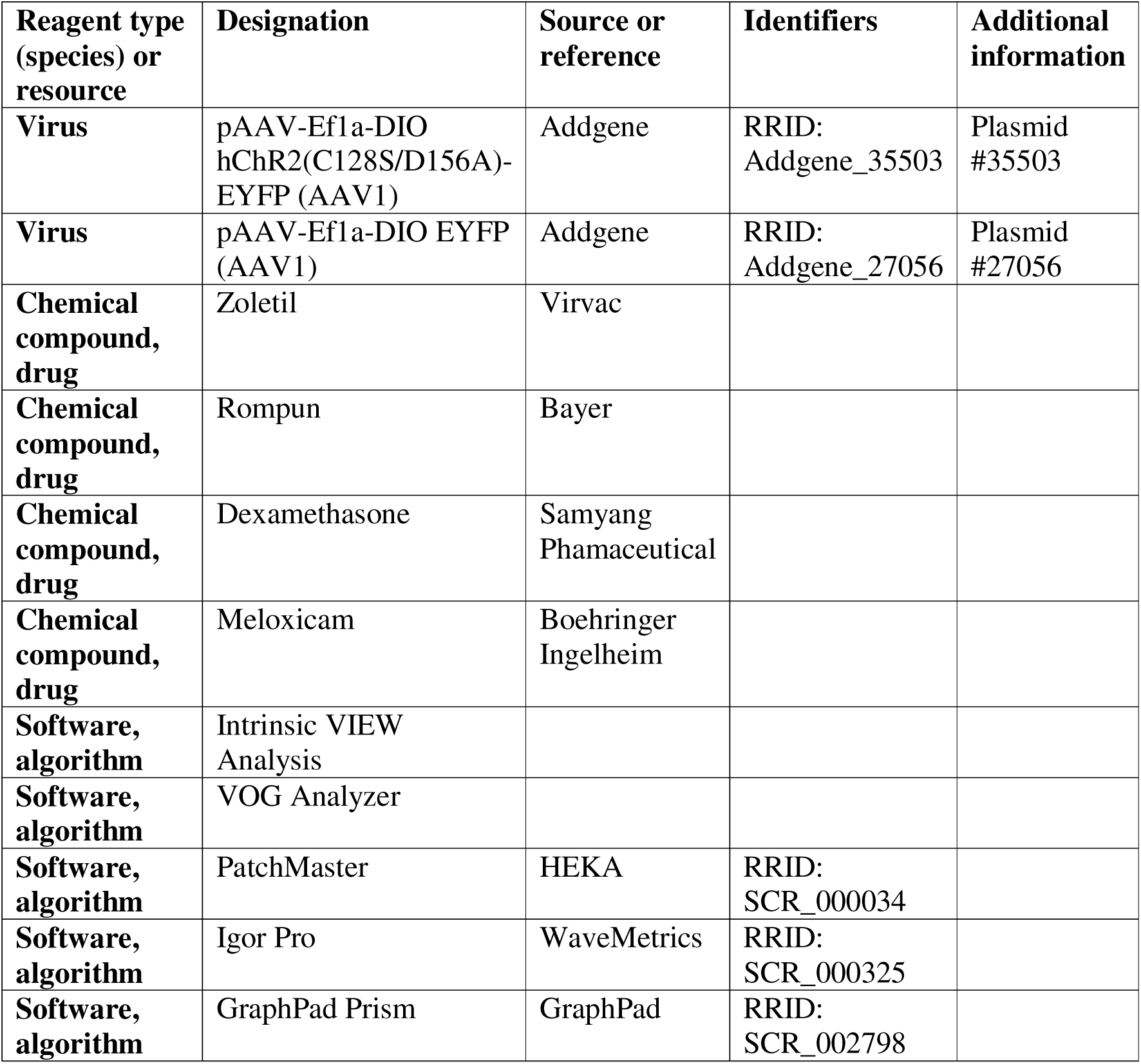

| Reagent type (species) or resource | Designation | Source or reference | Identifiers | Additional information |
| --- | --- | --- | --- | --- |
| **Virus** | pAAV-Ef1a-DIO hChR2(C128S/D156A)-EYFP (AAV1) | Addgene | RRID: Addgene_35503 | Plasmid #35503 |
| **Virus** | pAAV-Ef1a-DIO EYFP (AAV1) | Addgene | RRID: Addgene_27056 | Plasmid #27056 |
| **Chemical compound, drug** | Zoletil | Virvac |  |  |
| **Chemical compound, drug** | Rompun | Bayer |  |  |
| **Chemical compound, drug** | Dexamethasone | Samyang Phamaceutical |  |  |
| **Chemical compound, drug** | Meloxicam | Boehringer Ingelheim |  |  |
| **Software, algorithm** | Intrinsic VIEW Analysis |  |  |  |
| **Software, algorithm** | VOG Analyzer |  |  |  |
| **Software, algorithm** | PatchMaster | HEKA | RRID: SCR_000034 |  |
| **Software, algorithm** | Igor Pro | WaveMetrics | RRID: SCR_000325 |  |
| **Software, algorithm** | GraphPad Prism | GraphPad | RRID: SCR_002798 |  |

### Animals and stereotaxic surgery for virus injection and optic cannula implantation

The experimental procedures were approved by the Seoul National University Institutional Animal Care and Use Committee and performed the guidelines of the National Institutes of Health. In this study, 6- to 10-week-old *PCP2-Cre* mice (Jackson Laboratory, ME, USA), a driver line with high selectivity for *Cre* activity in PCs, were anesthetized with intraperitoneal injections of Zoletil/Rompun mixture (30 mg/kg and 10 mg/kg, respectively) and then they were placed on a stereotaxic frame (Narishige, Tokyo, Japan) for virus injection. AAV-Ef1a-DIO hChR2(C128S/D156A)-EYFP (SSFO) and AAV-EF1A-DIO EYFP were manually injected at a rate of 3 nl/ sec into the bilateral flocculi using *Picopump*. The glass pipette was set at a 34° angle and the targeted coordinates for the virus injection were at 5.48 mm dorsoventral (DV) and ±0.10 mm mediolateral (ML). After the mice were fully recovered (approximately 1 week) from the first surgery of virus injection, the second operation of optic cannula implantation (Ø1.25 mm Ceramic Ferrule, L=3.0 mm) near the cerebellar flocculus and head fixation were conducted. Unlike the fiber optic cannula, the head fixation components were hand-made (personalized design and production). Prior to the surgery, the mice were anesthetized using intraperitoneal injection of a mixture of Zoletil 50 (Virbac, 15mg/kg) and xylazine (Rompun, Bayer, 15mg/kg). Again, the mice were placed on placed on a stereotaxic frame (Narishige, Tokyo, Japan), and optic cannulae were implanted on both sides of the flocculi at ±2.80 mm ML.

### Confirmation of viral expression

After the completion of all experiments, sampling was performed using cardiac perfusion. The brain was then extracted and coronal sections were made at 30 µm intervals. Images were acquired and processed using a confocal microscope (Zeiss LSM 7 MP, Carl Zeiss, Jena, Germany) and Zen software (Zeiss). The anatomical location of virus expression was confirmed by comparison with Paxinos mouse brain atlas(Keith B. J. Franklin, 2013).

### Slice preparation and electrophysiology recording

A acute brain slice preparation and electrophysiological experiments were performed as previously described (Kim & Kim, 2020). Here, 5 to 9-week-old mice were anesthetized using isoflurane and decapitated. To acquire the records of floccular PCs, 250 μm thick coronal slices of the cerebellar flocculus were obtained from mice using a vibratome (VT1200S, Leica). However, to acquire FTNs data, 200 μm thick coronal slices of the cerebellar MVN were obtained. The ice-cold cutting solution contained 75 mM sucrose, 75 mM of NaCl, 2.5 mM of KCl, 7 mM of MgCl2, 0.5 mM of CaCl2, 1.25 mM of NaH2PO4, 26 mM of NaHCO3, and 25 mM of glucose, with bubbled 95% O2 and 5% CO2. The slices were immediately transferred to the artificial cerebrospinal fluid (ACSF) containing 125 Mm of NaCl, 2.5 Mm of KCl, 1 mM of MgCl2, 2 mM of CaCl2, 1.25 mM of NaH2PO4, 26 mM of NaHCO3, and 10 mM of glucose, with bubbled 95 % O2 and 5 % CO2. Subsequently, they were recovered at 32 °C for 30 min and at room temperature for 1 hr. Brain slices were placed in a submerged chamber perfused with the ACSF for at least 10 min before recording. Whole-cell recordings were performed at 29.5–30 °C. We used recording pipettes (3–4 MΩ) filled with 9 mM of KCl, 10 mM of KOH, 120 mM of K-gluconate, 3.48 mM of MgCl2, 10 mM of HEPES, 4 mM of NaCl, 4 mM of Na2ATP, 0.4 mM of Na3GTP, and 17.5 mM of sucrose (pH 7.25). Electrophysiological data were acquired using an EPC9 patch-clamp amplifier (HEKA Elektronik) and PatchMaster software (HEKA Elektronik) at a sampling frequency of 20 kHz, and the signals were filtered at 2 kHz. All electrophysiological recordings of the PCs were acquired in the flocculus, which is a lobe-like structure of the cerebellum. The spontaneous firing rate was analyzed using Igor Pro (WaveMetrics) and normalized to the average firing rate during the baseline period. Whole-cell recordings of FTNs were acquired from the MVN that exhibited EYFP-expressing *boutons en passant* from floccular PCs. Inhibitory synaptic inputs were completely blocked using 100 mM of picrotoxin (Sigma Millipore) during PC recordings, and strychnine (1 mM) was added to block the glycinergic input in FTN recordings. Patch pipettes (3–4 MΩ) were pulled from borosilicate glass and filled with an internal solution (pH 7.25) containing 9 mM of KCl, 10 mM of KOH, 120 mM of K-gluconate, 3.48 mM of MgCl2, 10 mM of HEPES, 4 mM of NaCl, 4 mM of Na2ATP, 0.4 mM of Na3GTP, and 17.5 mM of sucrose for testing in vitro recordings, ex vivo PC excitability, and ex vivo FTN recordings. To examine the excitability of cerebellar PCs, we provided a series of square-wised current injections that ranged from 100–900 pA with increments of 100 pA for 500 ms. To investigate the excitability of FTNs in the MVN, we also provided current injections from -150 pA to 150 pA with increments of 50 pA for 1000 ms.

### Optokinetic reflex behavior test

First, three sessions of acclimation were conducted. During each session, the mouse was restrained using a custom-made restrainer for 30 min with and without light for habituation to the recording environment. Calibration was performed as described by Stahl et al. (2000). This allowed the conversion of the pupil into eye rotation (Stahl et al., 2000). Next, prior to oculomotor training, three basal oculomotor performances including OKR, VOR in the dark (dVOR) and VOR, in the light (lVOR), were measured. During the recording of eye movements, it was necessary to control the pupil dilation that was induced by the absence of light in the dark condition. Thus, physostigmine salicylate solution (eserine; Sigma Millipore) was given to mice under brief isoflurane anesthesia. The concentration of eserine solution was increased from 0.1%, 0.15% and 0.2% based on the pupil size. Basal oculomotor performances were described in gains. For the actual learning protocol, we adopted OKR, and it consisted of five training sessions (10 min per session), six checkup points and 24 h of consolidation period. During the training sessions, the mice were visually stimulated with a sinusoidally rotating drum (±5 degrees). Completely trained mice were placed back into their home cage, which was stored in the dark condition until the last gain checkup points. To assess the impact of optogenetic manipulation during the acquisition period, the training consisted of three sessions, each lasting 15 minutes.

### Optogenetic manipulation during optokinetic reflex

For in-vivo optogenetic manipulation, SSFO-expressing mice were stimulated by 473nm blue laser and 593nm yellow laser for activation and deactivation, respectively. Each optostimulation was briefly given for 2 seconds at 10.5 mW. During the behavior assessment to determine the critical time window of memory consolidation, both SSFO and EYFP-expressing mice were subjected to optogenetic manipulation during the OKR (optokinetic reflex) training. The training was conducted twice, with each manipulation occurring at a different time point. Following the first training session, a break period of 2 to 3 weeks was implemented to reset the learning conditions before the commencement of the second training.

### Gain analysis

Gains acquired from basal oculomotor performances and OKR learning calculated by a ratio of evoked eye movements to the movement of screen or turn-table as visual or vestibular stimulus, respectively. A custom-built LabView (National Instrument) analysis tool was used for all of the calculations.

### Statistical analysis

Graph plotting and statistical analysis were performed using GraphPad Prism (GraphPad Software Inc, CA, USA). The hypothesis was tested by using paired and unpaired t-tests between sample pairs using GraphPad Prism. Statistical significance was set at p< 0.05. Asterisks denoted in the graph indicate the statistical significance. * indicates p < 0.05, ** < 0.01, *** < 0.001, and **** < 0.0001. The test name and statistical values are presented in each figure legend. Data are shown as mean ± SEM and statistical evaluations were performed using, independent t test, one-way repeated-measured (RM) ANOVA and two-way RM ANOVA with post hoc Tukey’s test and Sidak’s test and were performed.

## Author Details

***Jewoo Seo***

Department of Physiology, Seoul National University College of Medicine, Seoul, Republic of Korea

Department of Biomedical Sciences, Seoul National University College of Medicine, Seoul, Republic of Korea

**Contribution:** Conceptualization, Resources, Data curation, Software, Formal analysis, Validation, Investigation, Visualization, Methodology, Writing – original draft, review and editing

**Competing interests:** No competing interests declared

***Yong Gyu Kim***

Department of Physiology, Seoul National University College of Medicine, Seoul, Republic of Korea

Department of Biomedical Sciences, Seoul National University College of Medicine, Seoul, Republic of Korea

**Contribution:** Conceptualization, Data curation, Software, Formal analysis, Investigation Methodology

**Competing interests:** No competing interests declared

***Yong-Seok Lee***

Department of Physiology, Seoul National University College of Medicine, Seoul, Republic of Korea

Department of Biomedical Sciences, Seoul National University College of Medicine, Seoul, Republic of Korea

Memory Network Medical Research Center, Neuroscience Research Institute, Wide River Institute of Immunology, Seoul National University College of Medicine, Seoul, Korea

**Contribution:** Conceptualization, Supervision, Writing – review and editing

**Competing interests:** No competing interests declared

***Sang Jeong Kim***

Department of Physiology, Seoul National University College of Medicine, Seoul, Republic of Korea

Department of Biomedical Sciences, Seoul National University College of Medicine, Seoul, Republic of Korea

Memory Network Medical Research Center, Neuroscience Research Institute, Wide River Institute of Immunology, Seoul National University College of Medicine, Seoul, Korea

**Contribution:** Conceptualization, Supervision, Funding acquisition, Writing – review and editing

**Competing interests:** No competing interests declared

## Acknowledgments

This work was supported by National Research Foundation of Korea (NRF) grants funded by the Korean Government (MSIT) (NRF-2018R1A5A2025964 and 2022M3E5E8017970 to S.J. Kim).

