## Supplementary material for "Cerebellar systems consolidation driven by the temporal dynamics of Purkinje cell excitability": Supple Figures and Supple Figure Legends

**Supplementary Figure legends**

**Supplementary Figure 1*.* Depression of neuronal IE is prevented in SSFO-expressing PCs following motor learning**

a. Raw traces of PC firing under 400 pA current injection at 0 min post-learning (Top, EYFP; bottom, SSFO)
b. Mean firing rate (Hz) is significantly lower in the EYFP group (n = 8) at 0 min post-learning than in the SSFO group (n = 6) (two-way analysis of variance [ANOVA] multiple with Sidak’s multiple comparisons, **** p < 0.0001).

The error bars indicate ±SEM.

**Supplementary Figure 2*.* SSFO-expressing mice exhibits intact memory consolidation without optogenetic manipulation**

a. Gain changes in the SSFO group without optostimulation (w/o optostim) (left) and SSFO with optostimulation (w/ optostim) (right) groups following learning are shown. A total of 10 SSFO-expressing mice were tested under two different conditions: with and without optostimulation. Regardless of the test conditions, all mice exhibited a significant increase in gain after learning (SSFO w/o optostim, **** p < 0.0001; SSFO w/ optostim, **** p < 0.0001; repeated measure [RM] one-way analysis of variance [ANOVA], Tukey’s multiple comparisons). After 24 hr of consolidation period, the SSFO w/o optostim group showed robust maintenance of long-term memory (p = 0.0922; RM one-way ANOVA Tukey’s multiple comparisons), whereas the SSFO w/ optostim group showed impaired consolidation (*** p = 0.0003; RM one-way ANOVA Tukey’s multiple comparisons).
b. Comparison of consolidation percentages between the SSFO w/ and w/o optostim groups. The consolidation percentage in the SSFO group w/o optostim is distinctly greater than that in the SSFO group w/o optostim (paired t-test, ** p = 0.0024).

The error bars indicate ±SEM.

**Supplementary Figure 3*.* Plasticity of intrinsic excitability of nonFlocculus-targeting neurons(nonFTNs) in the medial vestibular nucleus (MVN)**

a. Raw traces representing the excitability of EYFP-expressing nonFTNs at 100 pA and 150 pA current injections under learned (n = 7) and naïve (n = 7) conditions. The vertical scale bar represents 10 mV and the horizontal scale bar represents 200 ms.
b. Comparison of the intrinsic excitability of EYFP-expressing nonFTNs under naïve and learned conditions. The excitability of the learned (n = 6) and naïve (n = 7) groups is comparable. They exhibit no significant difference.
c. Raw traces representing the excitability of SSFO-expressing nonFTNs at 100 pA and 150 pA current injections under learned (n = 8) and naïve (n = 6) conditions. The vertical scale bar represents 10 mV and the horizontal scale bar represents 200 ms.
d. Comparison of the intrinsic excitability of SSFO-expressing nonFTNs under naïve and learned conditions. The excitability of learned (n = 8) and naïve (n = 7) groups is comparable. They exhibit no significant difference.
e. Comparisons of the mean firing rates between nonFTNs of the EYFP (n = 7) and SSFO (n = 8) groups after learning. The mean firing rates of both groups are comparable. They exhibit no significant difference.

The error bars indicate ±SEM.

**Supplementary Figure 1.**

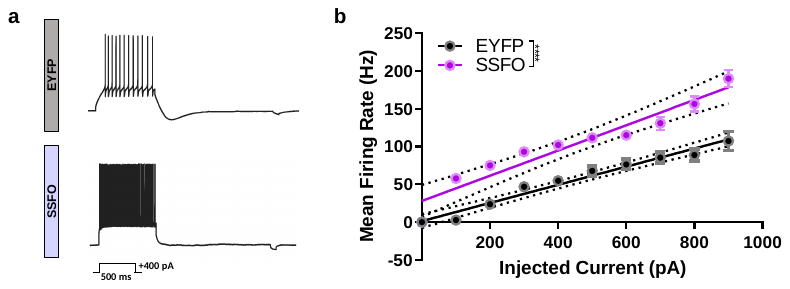

**Supplementary Figure 2.**

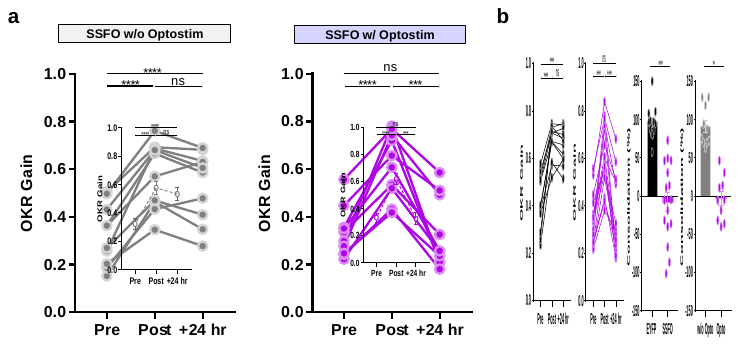

**Supplementary Figure 3**

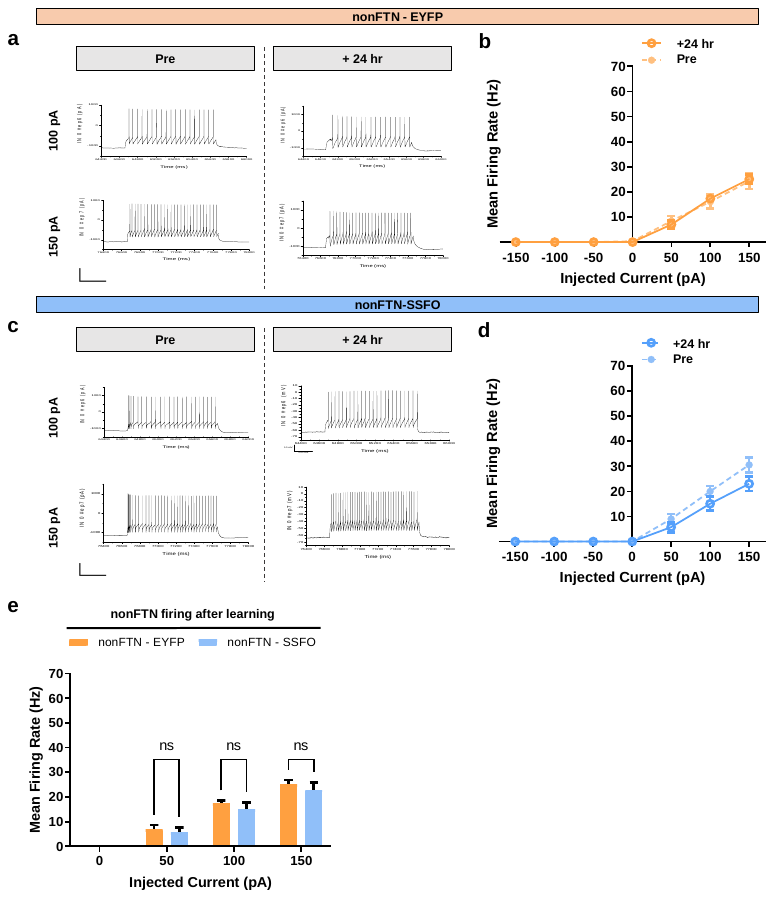
